# Occurrence and Antimicrobial Resistance Pattern of *E. coli* O157:H7 Isolated from Foods of Bovine Origin in Dessie and Kombolcha Towns, Ethiopia

**DOI:** 10.1101/2022.08.02.502434

**Authors:** Engidaw Abebe, Getachew Gugsa, Meselu Ahmed, Nesibu Awol, Yalew Tefera, Shimelis Abegaz, Tesfaye Sisay

## Abstract

*E. coli* are frequently isolated food-borne pathogens from meat, milk, and their products. Moreover, there has been a significant rise in the antimicrobial resistance patterns of *E. coli* O157:H7 to commonly used antibiotics. A cross-sectional study was conducted from October, 2019 to July, 2021 to estimate prevalence and identify associated factors of *E. coli* and *E. coli* O157:H7 and to determine antibiotic resistance pattern of *E. coli* O157:H7 from foods of bovine origin in Dessie and Kombolcha towns. A total of 384 samples were collected. Systematic and simple random sampling techniques were employed for sampling carcasses and milking cows, respectively. *E. coli* and *E. coli* O157:H7 were detected according to recommended bacteriological protocols. *E. coli* O157:H7 strains were evaluated for *in vitro* antimicrobial susceptibility using agar disk diffusion method. Both descriptive and inferential statistical techniques were applied to analyze the data. Overall prevalence rates of *E. coli* and *E. coli* O157:H7 were 54.7% and 6.5%, respectively. Highest prevalence rates of *E. coli* (79.6%) and *E. coli* O157:H7 (16.7%) were obtained from carcass swab and milk tank samples, respectively. Unlike *E. coli* O157:H7, a statistically significant difference in the *E. coli* prevalence (P<0.05) was observed among the different sample types. Multidrug resistance was observed among all isolates of *E. coli* O157:H7. All *E. coli* O157:H7 isolates (100.0%) were susceptible to Ampicillin, Sulfamethoxazole-trimethoprim, and Norfloxacin. On the contrary, all of the isolates (100%) were resistant to Penicillin G, Vancomycin, and Oxacillin. The current study indicated that different foods of bovine origin in the study area were unsafe for human consumption. Hence, good hygienic production methods should be employed to ensure the safety of foods of bovine origin.

## Introduction

Food-borne pathogens are the leading causes of human illness and death in the world [1]. Most microbial pathogens are zoonotic in nature and healthy food animals are reservoirs of many foodborne pathogens [2, 3]. In humans, consumption of foods of animal origin is a major source of exposure to food-borne pathogens [4]. Thus, people are at risk of being infected with pathogens from repository animals through the food chain [5].

Bacteria are the major cause of food-borne infections in humans [6]. Among different food-borne bacteria, *Escherichia coli* (*E. coli*) can get access to foods of animal origin from different sources [2], and these bacteria are frequently isolated food-borne pathogens from meat and meat products [7] and milk and dairy products [8].

*E. coli* are gram-negative, non-spore-forming, facultative anaerobic, and coliform bacteria belonging to the family *Enterobacteriaceae* that are residing in the intestines of animals and humans as normal microflora [3, 8-12]. The detection of *E. coli* in animal-derived foods is an indicator of fecal contamination and poor hygiene during production, storage, distribution, processing, or preparation of these food items, and the presence of other highly pathogenic microorganisms which can affect food safety and public health [13].

The species *E. coli* consists of a diverse and large group of bacteria [6]. Most *E. coli* strains are harmless [9]. However, some strains are pathogenic and can cause severe human illness [14]. Among these pathogenic strains, *E. coli* O157:H7 is one of the common and virulent food-borne bacterial pathogens [15] which is the subtype of Shiga toxin-producing *E. coli* strains [16]. This emerging food-borne bacterial strain is the leading cause of acute life-threatening infections such as hemolytic-uremic syndrome, hemorrhagic colitis, and thrombotic thrombocytopenic purpura in humans [1, 17, 18]. Cattle are the primary reservoirs of *E. coli* O157:H7 [3, 15, 18] and foods of bovine origin such as beef, milk, and dairy products are major sources and vehicles of human infection through the food chain [19].

Besides the magnitude, the increasing emergence and spread of antibiotic-resistant bacteria, multi-drug resistant zoonotic foodborne pathogens, in particular, have become a significant concern globally [20, 21]. The antimicrobial-resistant bacteria can be transmitted to humans through the food chain from food animal reservoirs [22]. Studies conducted in different areas indicated that there has been a significant rise in the antimicrobial resistance pattern of *E. coli* O157:H7 to commonly used antibiotics [23, 24].

Analysis of food to detect food-borne pathogens is essential to ensure food safety and to reduce and/or prevent the occurrence of food-borne infections in humans [25-27]. Particularly, the detection of food-borne pathogenic bacteria is critical for the control and prevention of some hazardous points in food production, processing, and/or distribution [28]. However, there is insufficient information related to the occurrence of food-borne infections in developing countries though the burden is high in those countries as compared to developed countries [15]. Despite there is growing tendency of reporting *E. coli* O157:H7 in beef and dairy products in recent times [1], only few studies have been reported related to the epidemiology and antibiotic resistance pattern of *E. coli* O157:H7 in Ethiopia [13, 15]. Furthermore, in most parts of Ethiopia, cow milk and beef are consumed as raw or undercooked which may prone people to pathogenic and drug-resistant food-borne bacteria. Hence, the objectives of the present study were to estimate the prevalence and identify associated factors of *E. coli* and *E. coli* O157:H7 and to determine the antibiotic resistance patterns of *E. coli* O157:H7 isolates from foods of bovine origin in Dessie and Kombolcha towns.

## Materials and Methods

### Ethics approval

This study was reviewed and approved by the Research Ethics Committee of the School of Veterinary Medicine, Wollo University. The study participants were informed about the study purpose.

### Study area

The study was conducted in Dessie and Kombolcha towns, South Wollo Zone, Eastern Amhara Region, Ethiopia (Fig 1). Dessie is the capital city of South Wollo zone which is located 401km to the northeast of Addis Ababa, the capital city of Ethiopia, and 480 km east of Bahir Dar, the capital city of Amhara Region [29]. The town is located at 11°8’N-11°46’ North latitude and 39°38’E-41013’East longitude. Topographically, Dessie town lies within an elevation ranges of 2,470 and 2,550 meters above sea level. It has a mean annual rainfall of 1100-1200 mm and the mean annual minimum and maximum temperatures of the town are 9°C and 23.7°C, respectively [30]. Administratively, Dessie town is subdivided into 18 urban and 8 rural Kebeles [31].

**Fig 1.**
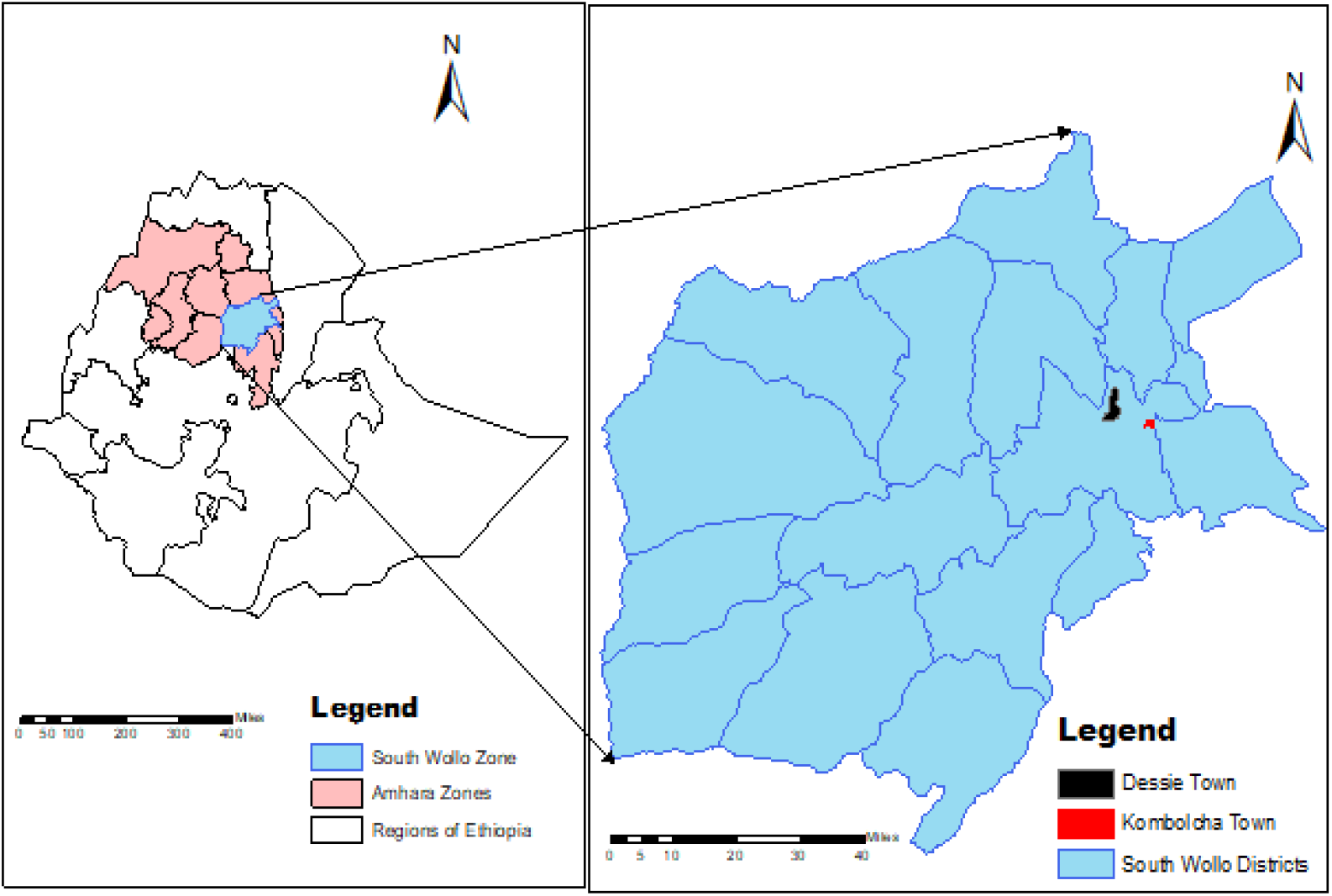
Map of the study areas.

Kombolcha is an industrial town situated at a distance of 376 km north of Addis Ababa, the capital city of Ethiopia, 23 km south-west of Dessie, the capital city of South Wollo zone, and 505 km from Bahirdar, the capital city of Amhara Region. The town is located at 11°6’ N latitude and 39°45’E longitudes with an elevation ranges from 1, 500 to 1, 840 meters above sea level [30]. The mean annual rainfall of Kombolcha town is 1046 mm and its annual minimum and maximum temperatures are 12.9°C and 28.1°C, respectively. Kombolcha town has a total of 11 administrative kebeles, 6 peri-urban and 5 urban [32].

### Study population

Udder and tank milk, milk product (yoghurt and cheese), carcass swab, and beef swab samples were collected from dairy farms, milk product shops, municipal, and ELFORA abattoirs, and butcher shops and restaurants in the study areas, respectively.

The total number of registered dairy farms in Kombolcha town at the time of sample collection was 164. In these farms, the total milking, dry and pregnant cows were 586, 266, and 386, respectively. The average daily milk yield in the town was 6,261 liters during the time of sample collection [33]. According to the document of Dessie Town Animal Production and Health Office [34], seven large-scale and well-organized dairy farms were found in Dessie town. With the exception of farmers having one to two milking cows, around 28 well-organized dairy farms were found in Dessie town through an assessment prior to sample collection. The total number of milking cows in those 28 farms in the town was around 196.

Averagely, 5 and 2 oxen were slaughtered per day in Dessie and Kombolcha municipal abattoirs, respectively. At Dessie municipal abattoir, 30 to 35 oxen were slaughtered on Friday of each week, and samples were collected on this day of every week. According to the information obtained from the meat inspector, the expansion of illegal field slaughtering practices and administrative problems are the main reasons for the low daily average number of oxen slaughtered at Kombolcha municipal abattoir, and samples at this abattoir were collected during Easter, Christmas, and Muslims’ holiday periods. There was no distinct demarcation of the slaughtering process in both municipal abattoirs: stunning, bleeding, skinning, evisceration, and carcass splitting areas. In Kombolcha municipal abattoir, there was no overhead rail and the whole slaughtering operation was carried out on the floor of a single room. The carcasses were hanged for splitting on overhead rail at Dessie municipal abattoir after bleeding, skinning, and evisceration procedures were done on the floor. The ELFORA abattoir, which is found in Kombolcha town, was equipped with slaughtering facilities, and slaughtering operation was carried out in rooms with clear division. At Kombolcha ELFORA abattoir, averagely 120 cattle were slaughtered per day.

### Study design

A cross-sectional study was conducted from October, 2019 to July, 2021 to estimate the prevalence of *E. coli* and *E. coli* O157:H7 and determine the antibiotic resistance pattern of *E. coli* O157:H7 from foods of bovine origin in Dessie and Kombolcha towns, Amhara, Ethiopia.

### Sample size determination

The sample size (n) was determined based on a statistical formula given by Thrusfield [35].

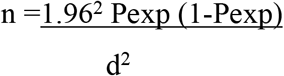

There was no previous published report related to the proportion of *E. coli* and *E. coli* O157:H7 from foods of bovine origin in Dessie and Kombolcha towns, hence an expected prevalence (Pexp) of 50% was used for sample size calculation with 95% confidence interval and 0.05 absolute precision (d). According to the above-given formula, the total sample size computed was 384.

After the assessment of the total number of sample sources (dairy farms, milk product shops, butcher shops, and restaurants, and the number of animals slaughtered at abattoirs) in the two study sites, the sample size of each sample type were allocated fairly. A total of 384 different samples of foods of bovine origin, including carcass swabs (n=162) from municipal and ELFORA abattoirs, udder milk (n=146) and milk tank (n=6) samples from dairy farms, yoghurt (n=36) and cheese (n=9) samples from milk product shops, and beef swabs (n=25) from butcher shops and restaurants were collected in the two selected study settings. With respect to the study site, 203 and 181 samples were collected from Kombolcha and Dessie towns, respectively.

### Sampling technique

A systematic random sampling method was employed to select carcass swab samples among cattle slaughtered at municipal and ELFORA abattoirs in study sites and every 3^rd^ cattle was selected. Milking cows from dairy farms in the study sites were selected using simple random sampling technique to collect udder milk samples. Moreover, tank milk, milk products (yoghurt and cheese), and beef swab samples were also collected randomly.

### Sample collection

Using sterile labeled screw cupped glass bottles, 25 ml of milk sample was collected from all quarters of selected individual milking cows in the middle of milking after the usual milking procedures were done by milkers in dairy farms. Tank milk samples, around 25 ml, were also collected from dairy farms using sterile labeled screw cupped glass bottles after the milk was mixed well. From milk product shops in study sites, approximately 25 ml/g yoghurt and cheese samples were collected aseptically using sterile labeled screw-capped glass bottles. At abattoirs, the carcass swab samples were collected using sterile cotton swabs from the surface and deep part of the selected carcass at five separate locations (neck, thorax, abdomen, breast, and crutch). The swab samples taken from different locations of the same carcass were pooled together and dipped into labeled test tubes containing 5 ml of sterile 0.85% NaCl solution. At butcher shops and restaurants, the beef swab samples were collected from different sites of the selected individual beef and placed into labeled test tubes containing 5 ml of sterile 0.85% NaCl solution. During the time of sample collection, all necessary data related to samples were recorded in a pre-designed format. The samples were shipped carefully on the day of collection using an ice box containing ice packs and processed within 24 hrs in Veterinary Microbiology Laboratory, School of Veterinary Medicine, Wollo University, Dessie, Ethiopia.

### Isolation and identification of *E. coli* and *E. coli* O157:H7

Detection of *E. coli* and *E. coli* O157: H7 in all collected samples was conducted according to Quinn et al. [36] bacteriological protocol for isolation and identification of these bacteria with a slight modification. The bacteriological media used for isolation and identification of these bacteria were prepared according to the instructions of the manufacturers. After each original sample was homogenized, 1 ml of the test sample was transferred into 9 ml sterile peptone water (Micromaster, India) and incubated aerobically at 37°C for 24 hrs. The pre-enriched samples were further inoculated into MacConkey broth (Blulux Laboratories Ltd., India) and incubated at 37°C for 24 hrs for selective enrichment. The enrichments were then streaked on MacConkey Agar plates (HiMedia Laboratories Pvt. Ltd., India) by four flame technique, and plates were incubated at 37°C for 24 hrs. Pink-colored colonies (Fig 2) were aseptically streaked on nutrient agar plates (HiMedia Laboratories Pvt.Ltd., India) and incubated at 37°C for 24 hrs. For microscopic observation, a pure colony was selected from nutrient agar plates and subjected to Gram staining as per procedures described by Merchant and Packer [37] to determine the shape, arrangement, and Gram reactions of the isolates. Gram-negative, pink-colored with rod-shaped appearance and arranged in single or in pairs were suspected as *E. coli*. A single isolated colony was picked and streaked on Eosin Methylene Blue Agar (EMB) medium (Sisco Research Laboratories Pvt. Ltd., India) and incubated aerobically at 37°C for 24 hrs. The presumptive *E. coli* colonies that showed greenish metallic sheen [38] (Fig 3) were picked up with a sterile inoculating loop and allowed to grow on nutrient agar plates (HiMedia Laboratories Pvt.Ltd., India) at 37ºC for 24 hrs for biochemical examination.

**Fig 2.**
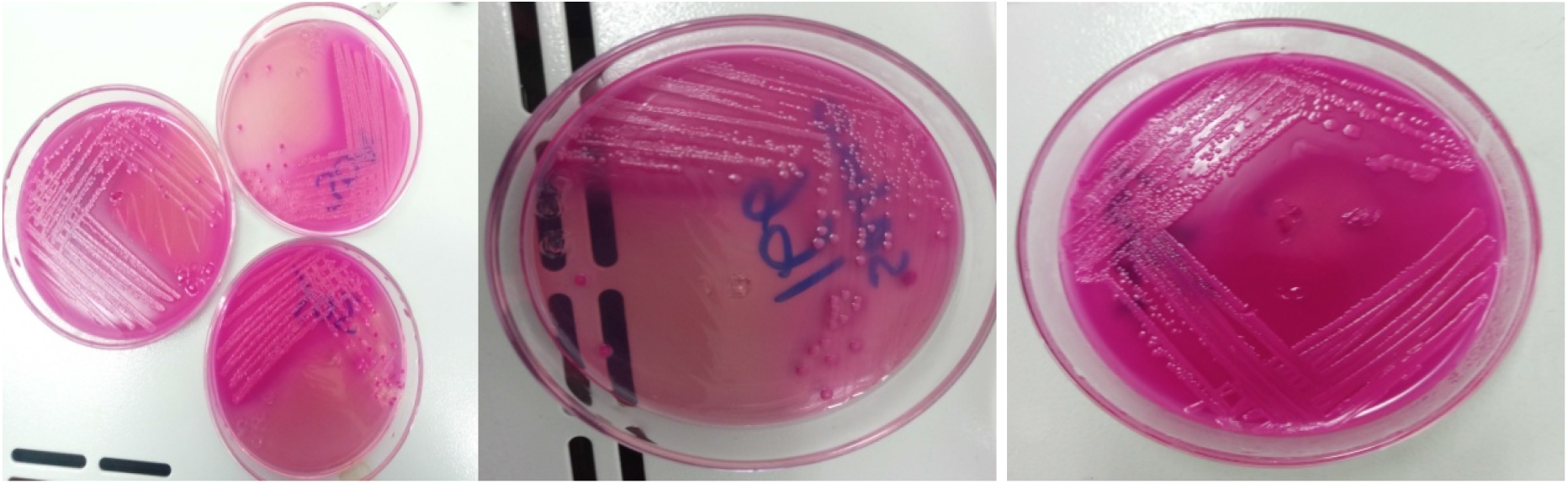
Growth on MacConkey agar plates.

**Fig 3.**
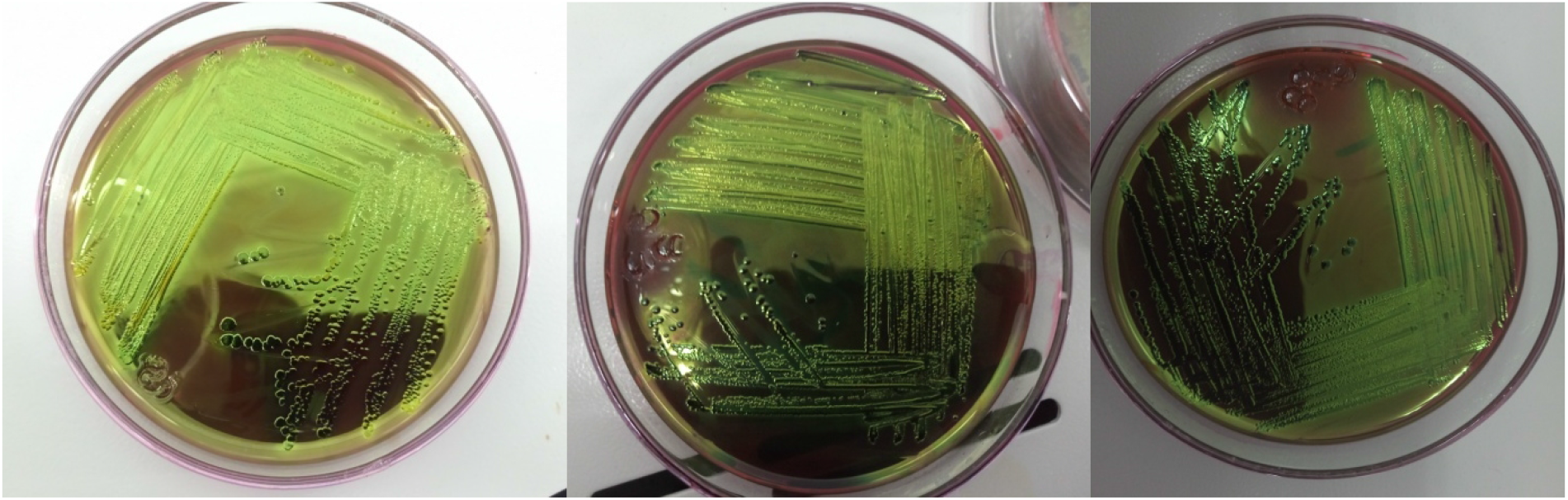
Growth on EMB agar plates.

Standard biochemical tests were used for confirmatory identification of the presumptive *E. coli* isolates [39, 40]. Slide catalase test of the isolates was performed according to MacFaddin [41] using 3% hydrogen peroxide (H_2_O_2_). Indole test was conducted according to Cheesbrough [42] by adding 0.5 ml of Kovac’s reagent (HiMedia Laboratories Pvt.Ltd., India) after the isolates were inoculated into Tryptone water broth (HiMedia Laboratories Pvt.Ltd., India) and incubated aerobically at 37ºC for 24 hrs. Methyl Red and Voges Proskauer tests were done according to Cheesbrough [42] after culturing the suspected colonies into MR-VP broth medium (Guangdong Huankai Microbial Sci.&Tech.Co., Ltd) at 37ºC for 24 hrs and adding 0.3 ml of 1% Methyl red solution (Dallul Pharmaceuticals Plc., Ethiopia), and 0.6 ml of 5% alpha naphthol (Loba Chemie Pvt.Ltd, Mumbai, India) and 0.2 ml of 40% potassium hydroxide solution (Unichem Laboratories Ltd., India), respectively. Citrate utilization test was performed according to the standard method of Simmons [43] by inoculating the colonies using stab and streak technique into Simmons citrate agar slants (HiMedia Laboratories Pvt.Ltd., India) and incubating at 37ºC for 24 hrs. Urease test for bacterial isolates was done according to the method Chakraborty et al. [44] by inoculating the isolates into Christensen Urea agar slant (Microxpress, India) and incubating at 37ºC for 24 hrs. According to Vanderzant and Splittstresser [45], TSI test was carried out by inoculating into TSI agar slants (Sisco Research Laboratories Pvt. Ltd, India) using stab and streak technique and incubated at 37ºC for 24 hrs. All biochemical tests were interpreted and isolates which were indole positive (pink to red color ring), methyl red positive (red color), Voges-Proskauer negative (no pink-red color at the surface), citrate negative (green butt and slant), urease negative (remaining yellow), and producing acid (both butt and slant changed to yellow color) with gas and without hydrogen sulfide production on TSI were confirmed to be *E. coli*.

The identified *E. coli* colonies were further subcultured onto SMAC agar plates (Guangdong Huankai Microbial Sci. & Tech. Co., Ltd., China) at 37ºC for 24 hrs to differentiate *E. coli* O157:H7 strain from other *E. coli* strains. Sorbitol-fermenters (pinkish colonies) were considered as non-O157:H7 *E. coli* strains whereas the non-sorbitol-fermenting isolates (colorless or pale colonies) were confirmed as *E. coli* O157: H7 strains (Fig 4).

**Fig 4.**
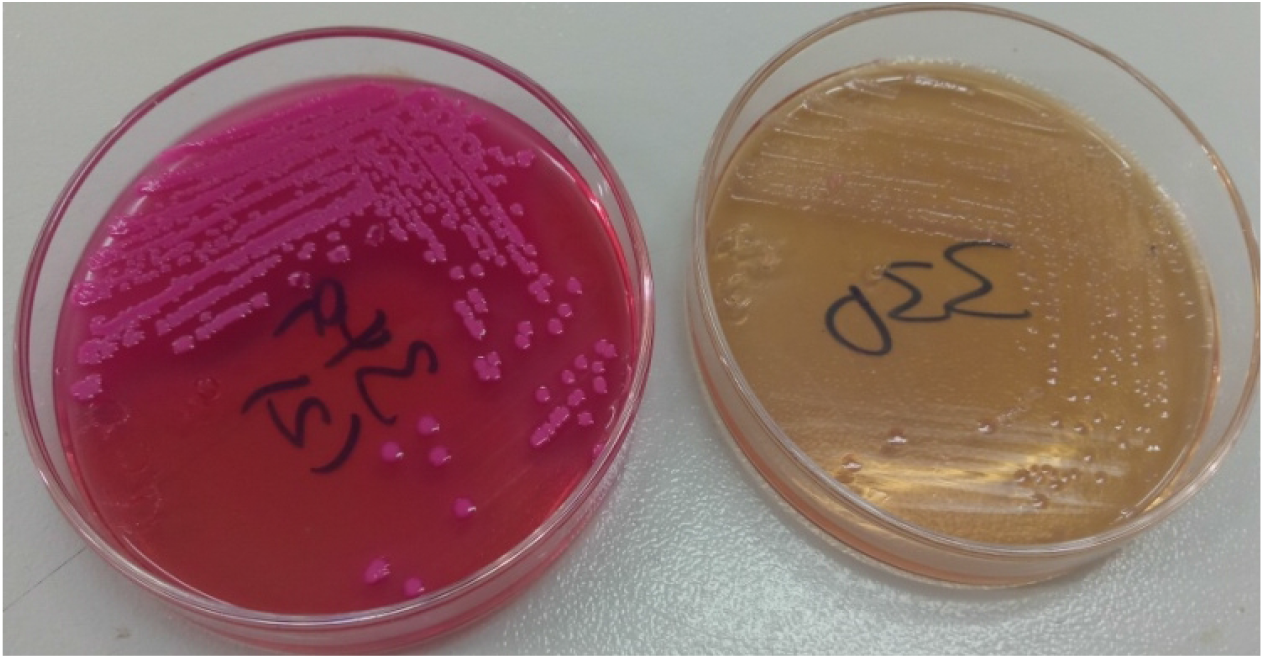
Growth on SMAC agar plates.

### Antimicrobial susceptibility testing of *E. coli* O157:H7

All *E. coli* O157:H7 isolates were evaluated for *in vitro* antimicrobial susceptibility using the agar disk diffusion method recommended by Bauer et al. [46]. The 16 antibiotic disks (Mast Group Ltd., Merseyside, U.K) and their concentrations used for the susceptibility testing were Amikacin (30μg), Amoxicillin (10µg), Ampicillin (25µg), Ceftriaxone (5μg), Ciprofloxacin (5μg), Doxycycline (30µg), Erythromycin (15µg), Gentamicin (10µg), Kanamycin (30µg), Nalidixic acid (30µg), Norfloxacin (2μg), Oxacillin (1μg), Penicillin G (10IU), Tetracycline (10µg), Sulphamethoxazole-trimethoprim (25µg), and Vancomycin (5μg).

*E. coli* O157:H7 isolates that had been confirmed biochemically were inoculated onto nutrient agar plates and incubated at 37°C for 24 hrs. After overnight incubation, colonies were transferred and diluted into test tubes containing 5 ml of sterile 0.85% saline solution and mixed thoroughly to generate a homogeneous suspension until the turbidity of the bacterial suspension achieved the 0.5 McFarland turbidity standards. A sterile cotton swab was dipped into the adjusted bacterial suspension and the excess inoculum was removed by lightly pressing the swab against the test tube’s upper inside wall.

To obtain uniform inoculums over the entire surface of the Mueller-Hinton agar plate (HiMedia Laboratories Pvt.Ltd., India), the swab containing the inoculum was spread evenly via a repeated rubbing procedure. The selected antibiotic-impregnated disks were placed at a minimum distance of 24 mm on the surface of the inoculated plate using sterile thumb forceps after the plates dried for 3 to 5 minutes and gently pressed with the point of a sterile forceps to ensure the complete contact between the disk and the agar surface. Within 15 minutes following the deposit of the disks, the plates were inverted and incubated at 37°C for 24 hrs. After 24 hrs of incubation, the zones of growth inhibition around each of the antibiotic disks were observed. The diameters of inhibition zones were measured using a digital caliper and the findings were recorded in a pre-designed format. The inhibition zone results around individual antibiotic disks were interpreted and the isolates were classified as Sensitive (S), Intermediate (I), and Resistant (R) according to the interpretation tables of the Clinical and Laboratory Standard Institute [47-50], Arabzadeh et al. [51], Reza et al. [52], Tadesse et al. [53], and TMCC [54].

### Standard organisms for quality control

To monitor the performance of the laboratory test and ensure accurate test results, the standard strains of *E. coli* obtained from Amhara Public Health Institute (APHI) Dessie branch, were used as control strains.

### Data management and analysis

All raw data collected from the study were summarized, compiled, entered, and coded in Microsoft Excel 2007 spreadsheet and imported to STATA Version 12 software for statistical analysis. Both descriptive and inferential statistical techniques were applied to analyze and present the different data types collected from the current study. Among descriptive statistics, frequency and/or percentage were calculated. Chi-square test (χ^2^) and binary logistic regression were computed to determine the association of different risk factors with contamination of *E. coli* and *E. coli* O157: H7 from different foods of bovine origin and the degree of association was determined using Odds ratio (OR) with 95% confidence interval (CI). Statistical significance was considered at P-value less than 0.05.

## Results

### Overall prevalence

Out of the total of 384 examined samples, 210 (54.7%) and 25 (6.5%) were *E. coli* and *E. coli* O157:H7 positive, respectively (Table 1).

**Table 1.** Prevalence of *E. coli* and *E. coli* O157:H7 among the sample types and study sites.

| Variables | No. of<br>examined | No. of positive (%) | | $\chi^2$-value | | P-value | |
| --- | --- | --- | --- | --- | --- | --- | --- |
|  |  | *E. coli* | *E. coli* | *E. coli* | *E. coli* | *E. coli* | *E. coli* |
|  |  |  | O157:H7 |  | O157:H7 |  | O157:H7 |
| Sample type |  |  |  |  |  |  |  |
| Udder milk | 146 | 63 (43.2) | 14 (9.6) | 82.871 | 6.926 | 0.000 | 0.226 |
| Tank milk | 6 | 2 (33.3) | 1 (16.7) |  |  |  |  |
| Yoghurt | 36 | 5 (13.9) | 0 (0.0) |  |  |  |  |
| Cheese | 9 | 1 (11.1) | 0 (0.0) |  |  |  |  |
| Beef swab | 25 | 10 (40.0) | 1 (4.0) |  |  |  |  |
| Carcass swab | 162 | 129 (79.6) | 9 (5.6) |  |  |  |  |
| Study site |  |  |  |  |  |  |  |
| Dessie | 181 | 95 (52.5) | 8 (4.4) | 0.6695 | 2.4584 | 0.413 | 0.117 |
| Kombolcha | 203 | 115 (56.7) | 17 (8.4) |  |  |  |  |
| Overall | 384 | 210 (54.7) | 25 (6.5) |  |  |  |  |

The sample type based prevalence of *E. coli* from carcass swab, udder milk, beef swab, tank milk, yoghurt, and cheese samples was 79.6%, 43.2%, 40.0%, 33.3%, 13.9%, and 11.1%, respectively. A statistically significant difference in the *E. coli* prevalence (P<0.05) was observed among the different sample types of foods of bovine origin. Among the examined sample types, the highest (16.7%) and lowest (0.0%) prevalence rates of *E. coli* O157:H7 were recorded from tank milk and milk products, respectively. The difference in the prevalence of *E. coli* O157:H7 among different sample types was not statistically significant (P>0.05) (Table 1).

With respect to the study site, the prevalence rates of *E. coli* (52.5% and 56.7%) and *E. coli* O157:H7 (4.4% and 8.4%) were found in Dessie and Kombolcha towns, respectively. There was no statistically significant difference in the prevalence rates of the isolates between the two study sites (P>0.05) (Table 1).

The odds of detection of *E. coli* were 31.27 times higher among carcass swab samples than in cheese samples and it was statistically significant (P<0.05) (Table 2).

**Table 2.** Bivariate logistic regression result of *E. coli* among different sample types.

| Sample type predictor | *E. coli* |  |
| --- | --- | --- |
|  | OR (95% CI) | P-value |
| Cheese | Reference |  |
| Tank milk | 4.0 (0.27-58.56) | 0.311 |
| Yoghurt | 1.29 (0.13-12.66) | 0.827 |
| Udder milk | 6.07 (0.74- 49.81) | 0.093 |
| Beef swab | 5.33 (0.57- 49.48) | 0.141 |
| Carcass swab | 31.27 (3.78-258.91) | 0.001 |

### Prevalence of *E. coli* and *E. coli* O157:H7 among variables of different sample types

The recorded prevalence rate of *E. coli* O157:H7 in milk samples from cows with treatment history was 13.7%. The difference in the prevalence of *E. coli* O157:H7 among treatment history categories was statistically significant (P<0.05). The prevalence of *E. coli* O157:H7 from milk samples was higher in Kombolcha town (14.0%) than in Dessie town (1.9%). The difference in the prevalence of *E. coli* O157:H7 between the two sites was statistically significant (P<0.05) (Table 3).

**Table 3.** Prevalence of *E. coli* and *E. coli* O157:H7 among different variables of milk samples.

| Variables | No. of<br>examined | No. of positive | | $\chi^2$-value | | P-value | |
| --- | --- | --- | --- | --- | --- | --- | --- |
|  |  | *E. coli* | *E. coli*<br>O157:H7 | *E. coli* | *E. coli*<br>O157:H7 | *E. coli* | *E. coli*<br>O157:H7 |
| Study site |  |  |  |  |  |  |  |
| Dessie | 52 | 20 (38.5) | 1 (1.9) | 0.598 | 5.610 | 0.440 | 0.018 |
| Kombolcha | 100 | 45 (45.0) | 14 (14.0) |  |  |  |  |
| Sample type |  |  |  |  |  |  |  |
| Udder milk | 146 | 63 (43.2) | 14 (9.6) | 0.227 | 0.325 | 0.634 | 0.569 |
| Tank milk | 6 | 2 (33.3) | 1 (16.7) |  |  |  |  |
| Farm System |  |  |  |  |  |  |  |
| Intensive | 131 | 56 (42.7) | 14 (10.7) | 0.000 | 0.714 | 0.993 | 0.398 |
| Semi Intensive | 21 | 9 (42.9) | 1 (4.8) |  |  |  |  |
| Treatment history |  |  |  |  |  |  |  |
| No | 50 | 18 (36.0) | 1 (2.0) | 1.392 | 5.186 | 0.238 | 0.023 |
| Yes | 102 | 47 (46.1) | 14 (13.7) |  |  |  |  |
| Milking practice |  |  |  |  |  |  |  |
| Excellent | 3 | 0 (0.0) | 0 (0.0) |  |  |  |  |
| Very good | 42 | 22 (52.4) | 4 (9.5) | 6.155 | 0.721 | 0.104 | 0.868 |
| Good | 104 | 43 (41.3) | 11 (10.6) |  |  |  |  |
| Poor | 3 | 0 (0.0) | 0 (0.0) |  |  |  |  |
| Farm hygiene |  |  |  |  |  |  |  |
| Excellent | 4 | 0 (0.0) | 0 (0.0) |  |  |  |  |
| Very good | 52 | 25 (48.1) | 6 (11.5) | 3.708 | 1.711 | 0.295 | 0.635 |
| Good | 85 | 35 (41.2) | 7 (8.2) |  |  |  |  |
| Poor | 11 | 5 (45.5) | 2 (18.2) |  |  |  |  |
| Total | 152 | 65 (42.8) | 15 (9.9) |  |  |  |  |

According to the result presented in Table 4, *E. coli* O157:H7 was not detected in milk products. The difference in the prevalence of *E. coli* among all hypothesized variables of milk products was not statistically significant (P>0.05).

**Table 4.** Prevalence of *E. coli* and *E. coli* O157:H7 among the variables of milk product samples.

| Variables | No. of<br>examined | No. of positive (%) | | $\chi^2$-value | P-value |
| --- | --- | --- | --- | --- | --- |
|  |  | *E. coli* | *E. coli* O15:7H7 | *E. coli* | *E. coli* |
| Study site |  |  |  |  |  |
| Dessie | 20 | 3 (15.0) | 0 (0.0) | 0.087 | 0.769 |
| Kombolcha | 25 | 3 (12.0) | 0 (0.0) |  |  |
| Sample Type |  |  |  |  |  |
| Yoghurt | 36 | 5 (13.9) | 0 (0.0) | 0.048 | 0.826 |
| Cheese | 9 | 1 (11.1) | 0 (0.0) |  |  |
| Equipment Type |  |  |  |  |  |
| Aluminum can | 2 | 0 (0.0) | 0 (0.0) | 0.322 | 0.570 |
| Plastic | 43 | 6 (14.0) | 0 (0.0) |  |  |
| Hygiene |  |  |  |  |  |
| Excellent | 6 | 0 (0.0) | 0 (0.0) | 4.350 | 0.114 |
| Very good | 20 | 5 (25.0) | 0 (0.0) |  |  |
| Good | 19 | 1 (5.3) | 0 (0.0) |  |  |
| Total | 45 | 6 (13.3) | 0 (0.0) |  |  |

There was no statistically significant difference in the prevalence of *E. coli* O157:H7 among all hypothesized variables of carcass swab samples (P>0.05). The prevalence of *E. coli* from carcass swab samples was higher in Kombolcha town (89.4%) than in Dessie town (72.9%) and the difference was statistically significant (P<0.05) (Table 5).

**Table 5.**
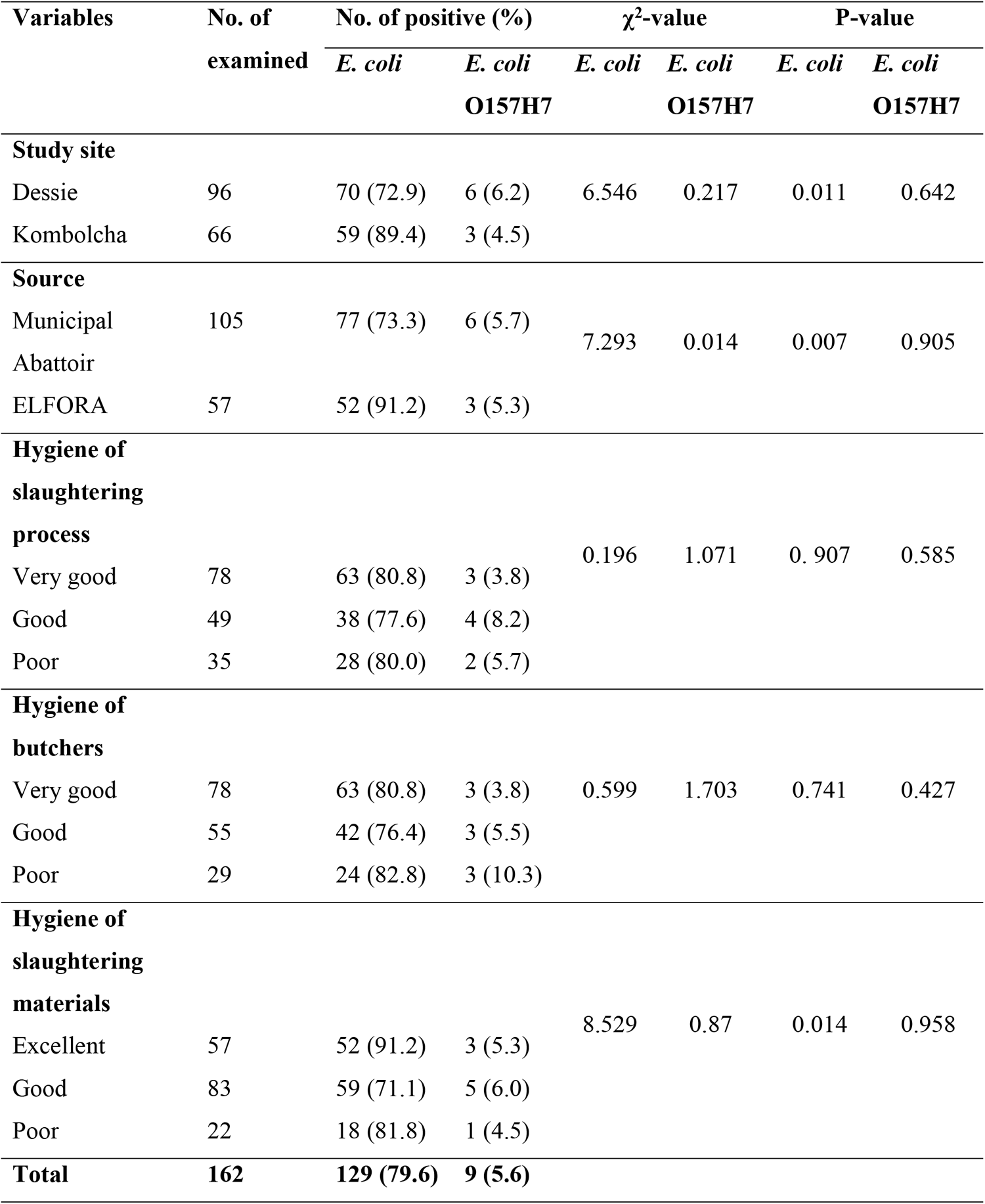
Prevalence of *E. coli* and *E. coli* O157:H7 among variables of carcass swab samples.

The proportion of *E. coli* in beef swab samples collected from butcher shops and restaurants in Kombolcha town (66.7%) was higher than in Dessie town (15.4%) and the difference was statistically significant (P<0.05). A higher prevalence of *E. coli* O157:H7 (50.0%) was obtained in beef swab samples collected from butcher shops having poor hygiene and the difference was statistically significant (P<0.05) as presented in Table 6.

**Table 6.**
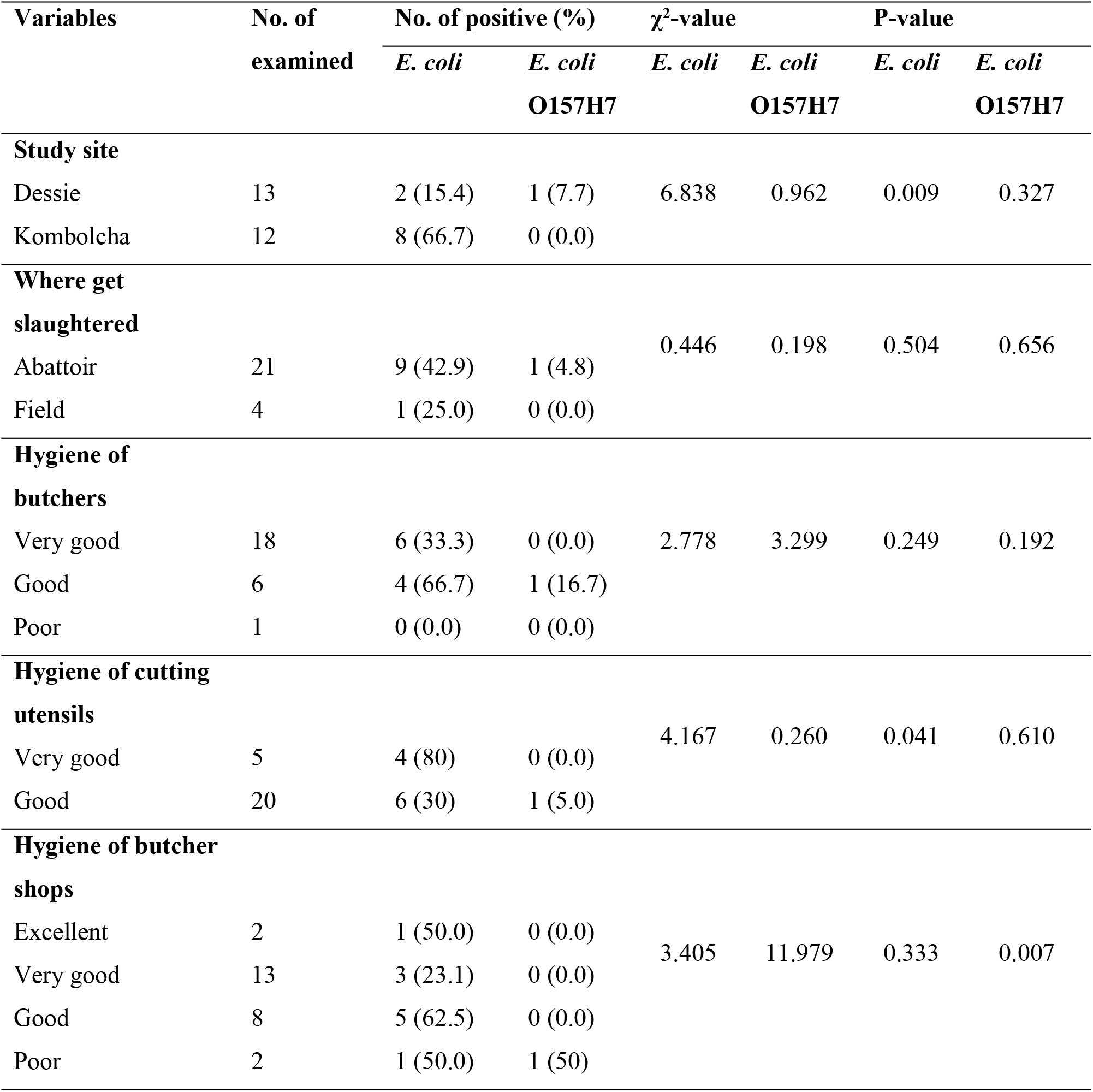
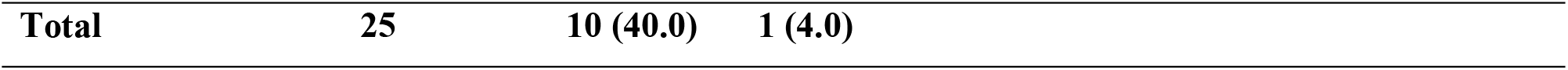
Prevalence of *E. coli* and *E. coli* O157:H7 among the variables of beef swab samples.

| Variables | No. of<br>examined | No. of positive (%) | | $\chi^2$-value | | P-value | |
| --- | --- | --- | --- | --- | --- | --- | --- |
|  |  | *E. coli* | *E. coli*<br>O157H7 | *E. coli* | *E. coli*<br>O157H7 | *E. coli* | *E. coli*<br>O157H7 |
| **Study site** |  |  |  |  |  |  |  |
| Dessie | 13 | 2 (15.4) | 1 (7.7) | 6.838 | 0.962 | 0.009 | 0.327 |
| Kombolcha | 12 | 8 (66.7) | 0 (0.0) |  |  |  |  |
| **Where get**<br>**slaughtered** |  |  |  |  |  |  |  |
| Abattoir | 21 | 9 (42.9) | 1 (4.8) | 0.446 | 0.198 | 0.504 | 0.656 |
| Field | 4 | 1 (25.0) | 0 (0.0) |  |  |  |  |
| **Hygiene of**<br>**butchers** |  |  |  |  |  |  |  |
| Very good | 18 | 6 (33.3) | 0 (0.0) | 2.778 | 3.299 | 0.249 | 0.192 |
| Good | 6 | 4 (66.7) | 1 (16.7) |  |  |  |  |
| Poor | 1 | 0 (0.0) | 0 (0.0) |  |  |  |  |
| **Hygiene of cutting**<br>**utensils** |  |  |  |  |  |  |  |
| Very good | 5 | 4 (80) | 0 (0.0) | 4.167 | 0.260 | 0.041 | 0.610 |
| Good | 20 | 6 (30) | 1 (5.0) |  |  |  |  |
| **Hygiene of butcher**<br>**shops** |  |  |  |  |  |  |  |
| Excellent | 2 | 1 (50.0) | 0 (0.0) | 3.405 | 11.979 | 0.333 | 0.007 |
| Very good | 13 | 3 (23.1) | 0 (0.0) |  |  |  |  |
| Good | 8 | 5 (62.5) | 0 (0.0) |  |  |  |  |
| Poor | 2 | 1 (50.0) | 1 (50) |  |  |  |  |

|  |  |  |  |
| --- | --- | --- | --- |
| **Total** | **25** | **10 (40.0)** | **1 (4.0)** |

### *In vitro* antimicrobial sensitivity pattern of *E. coli* O157:H7 isolates

The result of the *in vitro* antimicrobial sensitivity assay of the 25 *E. coli* O157:H7 isolates to the sixteen selected antimicrobial agents revealed that all strains (100.0%) were susceptible to Ampicillin, Sulfamethoxazole-trimethoprim, and Norfloxacin. On the contrary, all of the isolates (100%) were resistant to Penicillin G, Vancomycin, and Oxacillin. Moreover, high percentages of the isolates (92.0%) were also resistant to Erythromycin as presented in Table 7.

**Table 7.**
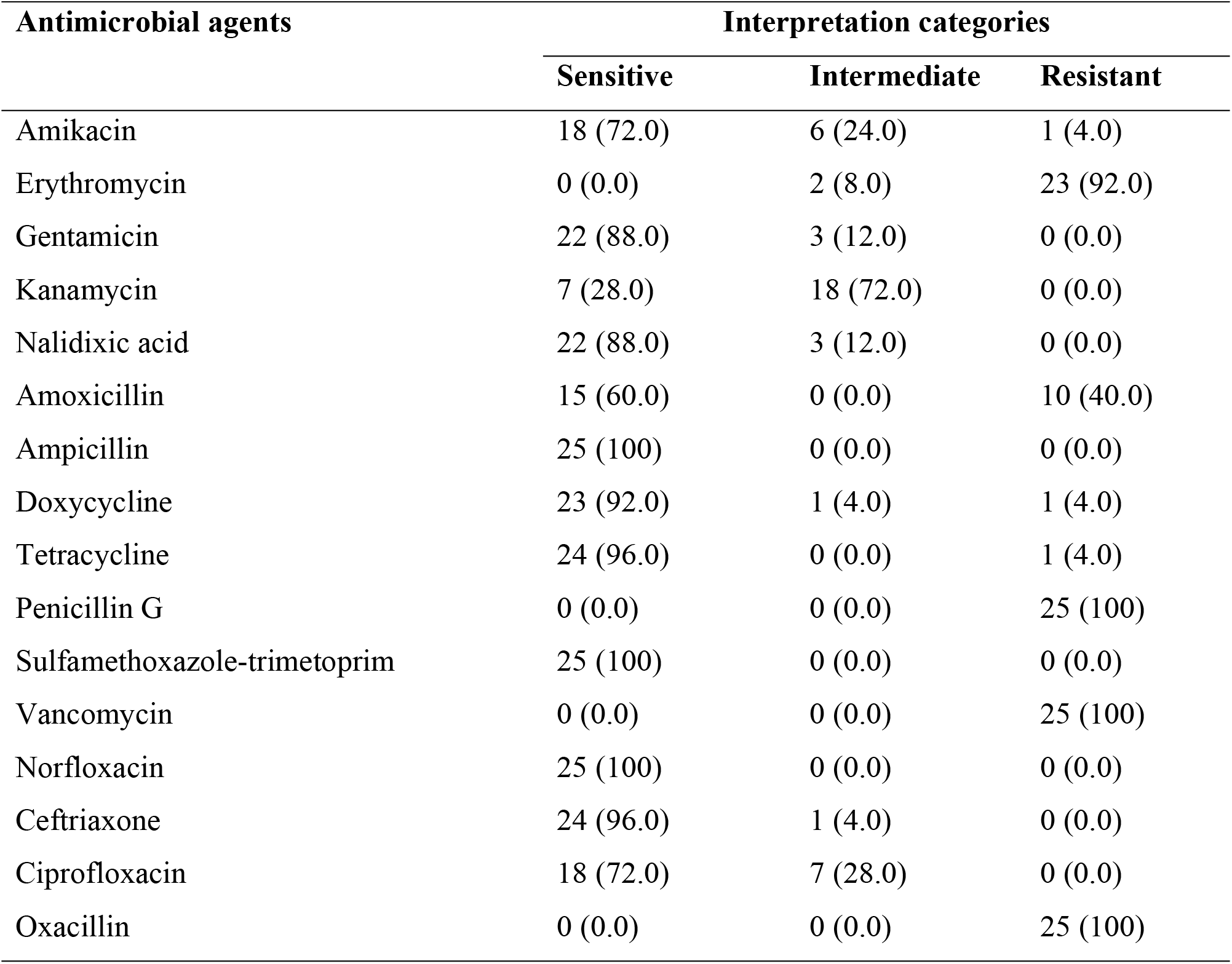
*In vitro* antimicrobial sensitivity pattern of *E. coli* O157:H7 isolated from different sample types of foods of bovine origin.

Multidrug resistance to more than three drugs was observed among all isolates of *E. coli* O157:H7. As shown in Fig 5, 14 (56.0%) and 11 (44.0%) of the isolates showed resistance to four and five drugs, respectively.

**Fig 5.**
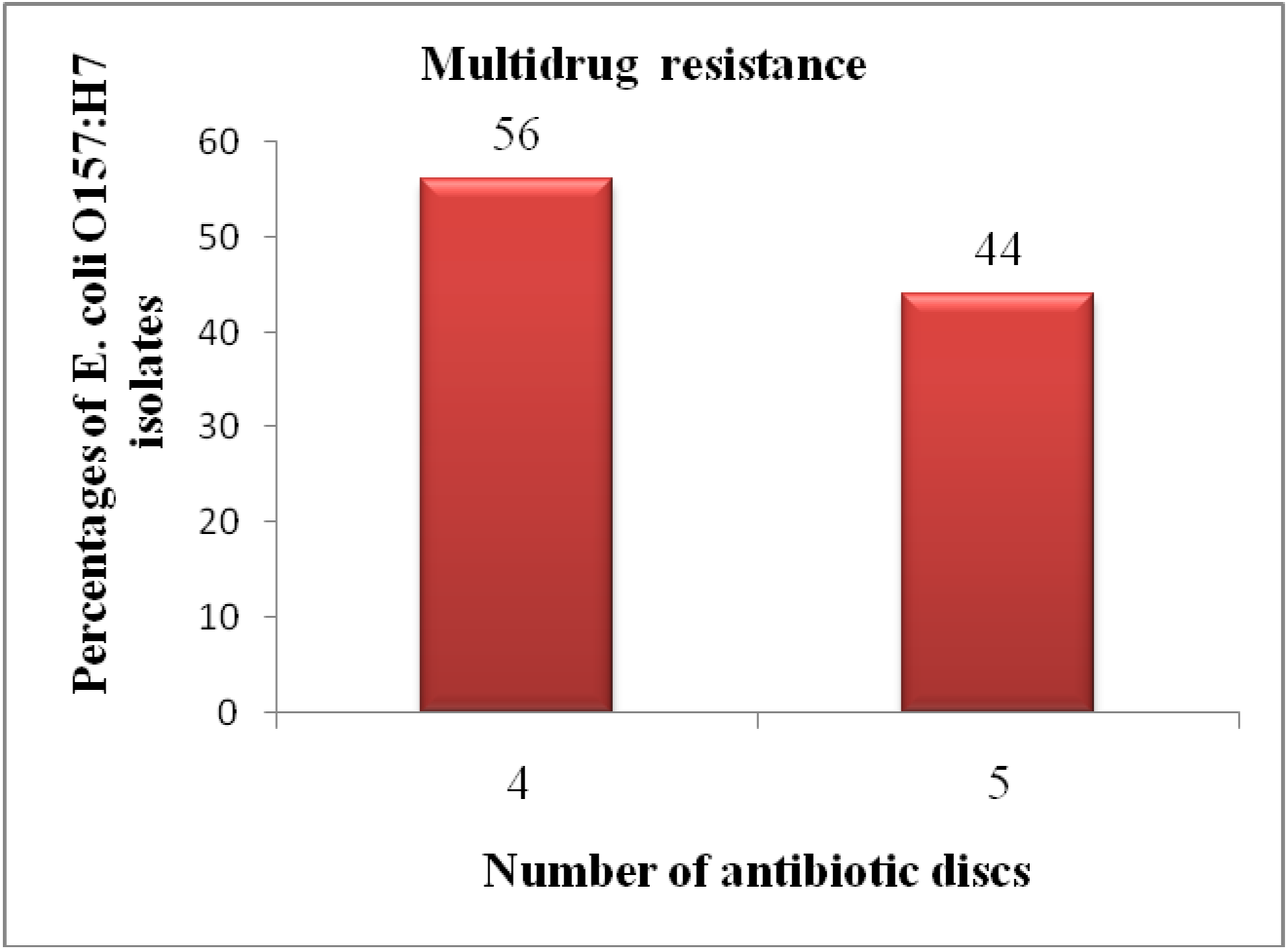
Multidrug resistance pattern of *E. coli* O157:H7 isolates.

## Discussion

The present study revealed an overall *E. coli* prevalence of 54.7% from different foods of bovine origin collected from different sources in the study areas. This prevalence was in agreement with previous studies reported by Limbu et al. [55] (55.0%), Soomro et al.) [56] (55.0%), Atsbha et al. [57] (57.29%), Reta et al. [58] (58.0%), Tadesse et al. [53] (51.2%), and Meshref [59] (52.6%) in Dharan (Nepal), Tandojam (Pakistan), Mekelle town, Jigjiga city, Mekelle town, and Beni-Suef governorate (Egypt), respectively.

However, in comparison to the present study, higher prevalence rates of *E. coli* were reported by Salauddin et al. [60] (100.0%), Baz et al. [61] (96.0%), Arjyal et al. [62] (92.0%), Balcha et al. [63] (62.5%), Gundogan and Avci [64] (74.0%), Lingathurai and Vellathurai [65] (70.0%), Altalhi and Hassan [66] (66.0%), Chyea et al. [67] (64.5%), and Ali and Abdelgadir [68] (63.0%) in Rangpur Division (Bangladesh), Kars city (Turkey), Kathmandu Valley (Nepal), Mekelle, Turkey, Madurai (South India), Taif region (Western Saudi Arabia), Malaysia, and Khartoum state, respectively.

On the other hand, the prevalence of *E. coli* in the current study was higher than the reports of Messele et al. [7] in Addis Ababa and Bishoftu towns (5.5%), Messele et al. [12] in central Ethiopia (Sebeta, Burayu, and Holeta towns) (7.1%), Kumar and Prasad [69] in and around Pantnagar (India) (8.14%), Yakubu et al. [70] in Sokoto Metropolis (Nigeria) (9.23%), Mengistu et al. [71] in Eastern Ethiopia (12.41%), Ngaywa et al. [72] in Kenya (13.8%), Mohammed et al. [73] in Dire Dawa city (15.89%), Ababu et al. [17] in Holeta District (19.0%), Hiwot et al. [74] in Arsi and East Shewa Zones (19.8%), Bedasa et al. [19] in Bishoftu town (20.0%), Sebsibe and Asfaw [75] in Jimma town (20.2%), Tadese et al. [76] in Ambo town (23.4%), Abebe et al. [77] in selected districts of Tigray (23.7%), Abayneh et al. [78] in Jimma town (23.9%), Yohannes [79] in Mekelle town (25.0%), Haileselassie et al. [80] in Mekelle city (27.3%), Hiko et al. [81] in Addis Ababa (29.0%), Momtaz et al. [82] in Iran (29.7%), Taye et al. [83] in Haramaya University abattoir (30.97%), Disassa et al. [8] in and around Asosa town (33.9%), Tadesse et al. [53] in Mekelle town (36.63%), Thaker et al. [84] in Anand Gujarat (India) (38.0%), Zerabruk et al. [85] in Addis Ababa (43.75%), Sobeih et al. [86] in Kafr El-Sheikh Governorate (Egypt) (44.44%), and Welde et al. [87] in and around Modjo town (46.26%).

The result obtained from the current bacteriological study revealed that the overall prevalence of *E. coli* O157:H7 was 6.5%. This finding was consistent with the previous reports of Gutema et al. [88] (6.3%), Ababu et al. [17] (5.2%), Beyi et al. [89] (4.5%), Reuben and Owuna [90] (4.5%), Sebsibe and Asfaw [75] (5.4%), Hiko et al. [91] (8.0%), Rahimi et al. [92] (8.2%), Vanitha et al. [93] (8.8%), and Tadese et al. [76] (9.1%) in Bishoftu town, Holeta District, central Ethiopia, Nasarawa State (Nigeria), Jimma town, Debre-Zeit and Modjo towns, Fars and Khuzestan provinces (Iran), Kerala (India), and Ambo town, respectively. However, the result found in the present study was higher than Dadi et al. [94] (0.0%) in Sebeta town (Ethiopia), Baz et al. [61] in Kars city (Turkey) (0.0%), Swai and Schoonman [95] in Tanga region (Tanzania) (0.0%), Abdissa et al. [15] in Addis Ababa and Debre Berhan cities (0.8%), Yakubu et al. [70] in Sokoto Metropolis (Nigeria) (1.92%), Mengistu et al. [71] in Eastern Ethiopia (2.06%), Geresu and Regassa [96] in the selected study settings of Arsi Zone (2.1%), Atnafie et al. [13] in Hawassa town (2.33%), Meshref [59] in Beni-Suef governorate (Egypt) (2.6%), Taye et al. [83] in Haramaya University abattoir (2.65%), Disassa et al. [8] in and around Asosa town (2.9%), Carney et al. [97] in Ireland (3.0%), Mcevoy et al. [98] in Ireland (3.2%), Ahmed and Shimamoto [99] in Egypt (3.4%), and Bedasa et al. [19] in Bishoftu town (3.5%).

On the other hand, the prevalence of *E. coli* O157:H7 found in this study was lower than the reports of Lingathurai and Vellathurai [65] (65.0%), Islam et al. [100] (52.4%), Llorente et al. [101] (36.1%), Chyea et al. [67] (33.5%), Bekele et al. [10] (13.3%), Hamid et al. [102] (12.0%), Balcha et al. [63] (11.3%), and Abebe et al. [77] (10.4%) in Madurai (South India), Bangladesh, Buenos Aires (Argentina), Malaysia, Addis Ababa, Addis Ababa, in and around Mekelle, and selected districts of Tigray, respectively. Such variations in *E. coli* and *E. coli* O157:H7 prevalence rates between present and other previous studies might be due to differences in management and hygienic practices in dairy and beef farms, standards and furnishings of abattoir and dairy farms, dairy cow herd health status (these bacteria are commonly isolated from mastitic milk), hygienic conditions in slaughterhouses and milking premises, cleanliness of milking and slaughtering utensils, hygienic practices during milking and slaughtering, water quality and its availability, and hygienic conditions of foods of bovine origin during handling, transportation, storage, and distribution up to consumption. Moreover, the variations could also arise from differences in study methods employed by researchers including sample source, sample size, sampling techniques, sample type, and methods of detection in laboratories.

The current study showed that the prevalence of *E. coli* was highest in carcass swab samples (79.6%) followed by udder milk (43.2%), beef swab (40.0%), tank milk (33.3%), yoghurt (13.9%), and cheese (11.1%). Unlike *E. coli* O157:H7, a statistically significant difference in the *E. coli* prevalence (P<0.05) was observed among different sample types of foods of bovine origin. The odds of detection of *E. coli* were 31.27 times higher among carcass swab samples than in cheese samples and it was statistically significant (P<0.05). At abattoirs, sanitation and hygiene are the crucial factors that contribute to meat contamination [103]. Poor hygienic practices at abattoirs during bleeding, skinning, evisceration, carcass washing, and splitting might be responsible for the contamination and higher magnitude of *E. coli* in carcass samples. In addition, the prevalence of *E. coli* O157:H7 in tank milk, udder milk, carcass swab, beef swab, yoghurt, and cheese samples was 16.7%, 9.6%, 5.6%, 4.0%, 0.0%, and 0.0%, respectively. The presence of *E. coli* in milk is not only regarded as faecal contamination but also an indicator of poor hygiene and sanitary practices during milking and further handling [8, 64, 66]. The higher proportion of *E. coli* O157:H7 in tank milk could be from different sources including unhygienic milking practices, cows infected with mastitis, milk handlers with poor hygiene, poor quality water, and inappropriately cleaned milk filtering utensils and tanks.

The prevalence of *E. coli* O157:H7 from milk samples was higher in Kombolcha town (14.0%) than in Dessie town (1.9%). The statistically significant difference (P<0.05) in the prevalence of *E. coli* O157:H7 among the two study sites could be associated with variation in hygienic practices in the dairy environment and herd health status of dairy farms. A higher prevalence of *E. coli* O157:H7 was recorded in milk samples from cows with teat treatment history (13.7%) than non treated cows (2.0%) and the difference was statistically significant (P<0.05). Cows with previous mastitis history are more likely to become infected than those which had never been exposed as they might remain in a carrier state as well as the ineffectiveness of mastitis treatment medicines [104]. The most common serotypes of *E. coli* recovered from mastitic milk are O157, O55, O111, O124, O119, O114, O26, and O44 [105]. Thus, the relatively high magnitude of *E. coli* O157:H7 in milk samples from cows with treatment history might be associated with environmental bovine mastitis.

In the present study, *E. coli* O157:H7 was not detected in milk products. According to Rahimi et al. [106], the survival of *E. coli* O157:H7 in foods is dependent on the acidity of the sample; when the pH falls below 3.5, the bacteria die. Thus, the absence of *E. coli* O157:H7 in yogurt and cheese samples in this study might be due to the acidity of these products and the temperature used during the processing of cheese.

The prevalence of *E. coli* from carcass swab samples was higher in Kombolcha town (89.4%) than in Dessie town (72.9%) and the difference was statistically significant (P<0.05). The variation could be due to the difference in hygienic practices at abattoirs. The proportion of *E. coli* in beef swab samples collected from butcher shops and restaurants in Kombolcha town (66.7%) was higher than in Dessie town (15.4%) and the difference was statistically significant (P<0.05). The variation could be due to the difference in hygienic practices during the slaughtering process at abattoirs and sanitation at butcher shops and restaurants. Moreover, a higher prevalence of *E. coli* O157:H7 (50.0%) was obtained in beef swab samples collected from butcher shops having poor hygiene and the difference was statistically significant (P<0.05). The higher occurrence of *E. coli* O157:H7 in beef swab samples collected from butcher shops having poor hygiene was not surprising since beef contamination is usually associated with poor hygiene.

The occurrence of antimicrobial resistance among foodborne pathogens is increasing [107]. The *E. coli* O157:H7 strains are heterogeneous with respect to antibiotic resistance [108]. The development of antimicrobial resistance in *E. coli* O157:H7 strains isolated from animals and humans [90] and the emergence of multidrug-resistant *E. coli* O157:H7 strains become a universal public health concern [109]. In the present study, multidrug resistance to more than three drugs was observed among all *E. coli* O157:H7 isolates. In brief, 56.0% and 44.0% of the isolates showed resistance to four and five drugs, respectively.

All isolates of *E. coli* O157:H7 (100%) were resistant to Penicillin G, Vancomycin, and Oxacillin. Moreover, high percentages of the isolates (92.0%) were also resistant to Erythromycin. The total resistance to Penicillin G was similar to the reports of Igbinosa and Chiadika [110] and Reuben et al. [111] who reported 100.0% resistance to Penicillin G in Benin City (Nigeria) and Nasarawa State (Nigeria), respectively. However, Msolo et al. [112] reported 85.0% resistance to Penicillin G in South Africa. The resistance of all isolates to Vancomycin was comparable to the report of Bedasa et al. [19] who reported 90.0% resistance to Vancomycin in Bishoftu town. The high frequency of resistance to Erythromycin was in agreement with the previous reports of Reuben and Owuna [90] in Nasarawa State, Nigeria, and Igbinosa and Chiadika [110] in Benin City (Nigeria) who reported 94.7% and 89.5% resistance to Erythromycin, respectively. The total resistance of the isolates to Oxacillin was higher than the report of Reuben and Owuna [90] who reported 84.2% resistance to Oxacillin.

On the contrary, all *E. coli* O157:H7 strains (100.0%) were susceptible to Ampicillin, Sulfamethoxazole-trimethoprim, and Norfloxacin. The total susceptibility to Sulfamethoxazole-trimethoprim was similar to previous findings of Tadese et al. [76], Bekele et al. [10], Beyi et al. [89], and Geresu and Regassa [96] who reported 100.0% sensitivity to Sulfamethoxazole-trimethoprim in Ambo town, Addis Ababa, central Ethiopia, and selected study settings of Arsi Zone, respectively. The total susceptibility to Norfloxacin was similar to the previous finding of Tadese et al. [76] (100.0%) in Ambo town. The 100.0% susceptibility to Ampicillin was similar to the previous report of Osaili et al. [113] who reported 100.0% sensitivity to Ampicillin in Amman City, Jordan.

Higher percentages of the isolates were also sensitive to Doxycycline (92.0%), Tetracycline (96.0%), Ceftriaxone (96.0%), Gentamicin (88.0), Nalidixic acid (88.0%), Amikacin (72.0%), and Ciprofloxacin (72.0%). The sensitivity of isolates to Gentamicin was comparable with the report of Bedasa et al. [19] who reported 82.5% sensitivity to Gentamicin in Bishoftu town. The susceptibility to Amikacin was consistent with Msolo et al. [112] who reported 70.0% sensitivity to Amikacin in South Africa. The sensitivity to Ciprofloxacin was consistent with the reports of Bekele et al. [10] in Addis Ababa and Reuben and Owuna [90] in Nasarawa State, Nigeria who reported 76.5% and 78.9% sensitivity to Ciprofloxacin, respectively. The high sensitivity to Ceftriaxone was consistent with the reports of Bedasa et al. [19], Atnafie et al. [13], and Haile et al. [114] who reported 100% sensitive isolates to Ceftriaxone in Bishoftu, Hawassa, and Jimma towns, respectively. The 96.0% sensitivity to Tetracycline was consistent with Haile et al. [114] in Jimma, Bekele et al. [10] in Addis Ababa, Osaili et al. [113] in Amman City (Jordan), and Bedasa et al. [19] in Bishoftu town who reported 100%, 100.0%, 100.0% and 97.5% sensitivity to Tetracycline. However, Welde et al. [87] reported 77.8% resistance to Tetracycline in and around Modjo town. According to Mokgophi et al. [115] and Qamar et al. [116], the extensive, indiscriminate and injudicious use of antibiotics in both veterinary medicine and public health leads to genetic modification in most bacterial strains for evolving resistance and an increase in the prevalence of resistance among pathogens.

## Conclusion and Recommendations

The high magnitude of *E. coli* contamination and finding of multidrug-resistant *E. coli* O157:H7 in the current study indicated that different foods of bovine origin in the study area were unsafe for human consumption. The high *E. coli* contamination of these products might be due to unhygienic practices at different stages of farm to fork process, as these bacteria are hygiene indicators. The multidrug resistance pattern of all *E. coli* O157:H7 isolates might be due to the injudicious and extensive use of antibiotics in both veterinary medicine and public health. In addition, slaughtering of cattle on the floor at municipal abattoirs, illegal open field slaughtering practices, unsanitary milk production, and further handling, and the community’s consumption habit of raw animal products could expose humans in study sites to multidrug-resistant *E. coli* O157:H7 and other pathogenic food-borne bacterial pathogens. Hence, good hygienic production methods should be employed to ensure the safety of different foods of bovine origin. Microbiological guidelines mainly the HACCP system and standardized slaughtering operations should be followed to improve meat safety. The emergence and spread of antibiotic-resistant pathogens should be assessed regularly and rational use of antibiotics should be practiced. Moreover, further studies on serotyping and molecular characterization of *E. coli* O157:H7 should be done at the study sites.

## Acknowledgements

The authors would like to thank Wollo University Research and Community Service Vice President Office and Wollo University Post Graduate Directorate Office for financial funding. We would like to extend our acknowledgement to dairy farm owners and/or managers, farm attendants, milk product shop workers, restaurant managers, and owners, butchers, abattoir workers, veterinarians, animal health and production officers, and other inhabitants in the study sites for their voluntariness and cooperation during sample collection.

